# Relation Extraction Between Bacteria and Biotopes from Biomedical Texts with Attention Mechanisms and Domain-Specific Contextual Representations

**DOI:** 10.1101/686501

**Authors:** Amarin Jettakul, Duangdao Wichadakul, Peerapon Vateekul

## Abstract

The Bacteria Biotope (BB) task is biomedical relation extraction (RE) that aims to study the interaction between bacteria and their locations. This task is considered to pertain to fundamental knowledge in applied microbiology. Some previous investigations have used feature-based models; others have presented deep-learning-based models such as convolutional and recurrent neural networks used with the shortest dependency paths (SDPs). Although SDPs contain valuable and concise information, sections of significant information necessary to define bacterial location relationships are often neglected. In addition, the traditional word embedding used in previous studies may suffer from word ambiguation across linguistic contexts.

Here, we present a deep learning model for biomedical RE. The model incorporates feature combinations of SDPs and full sentences with various attention mechanisms. We also used pre-trained contextual representations based on domain-specific vocabularies. In order to assess the model’s robustness, we introduced a mean F1 score on many models using different random seeds. The experiments were conducted on the standard BB corpus in BioNLP-ST’16. Our experimental results revealed that the model performed better (in terms of both maximum and average F1 scores; 60.77% and 57.63%, respectively) compared with other existing models.

We demonstrated that our proposed contributions to this task can be used to extract rich lexical, syntactic, and semantic features that effectively boost the model’s performance. Moreover, we analyzed the trade-off between precision and recall in order to choose the proper cut-off to use in real-world applications.

## 1 Introduction

Due to the rapid development of computational and biological technology, the biomedical literature is expanding at an exponential rate [1]. This situation leads to difficulty manually extracting required information. In BioNLP-ST 2016, the Bacteria Biotope (BB) task [2] followed the general outline and goals of previous tasks defined in 2011 [3] and 2013 [4]. This task aims to investigate the interactions of bacteria and its biotope; habitats or geographical entity, from genetic, phylogenetic, and ecology perspectives. It involves the *Lives_in* relation, which is a mandatory relation between related arguments, the bacteria and the location where they live. Information pertaining to the habitats where bacteria live is particularly critical in applied microbiology fields such as food safety, health sciences, and waste processing [2, 3, 4]. An example relation between bacteria and their location in this task is shown in Figure 1.

**Figure 1:**
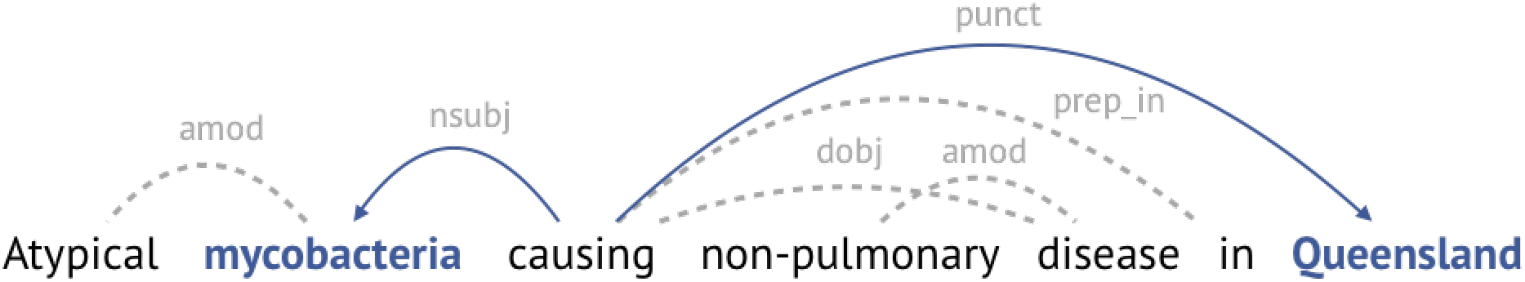
Example of the BB relation in a BB task. Bacteria “mycobacteria” and location “Queensland” are shown in blue, bold text. The dependencies are represented by arrows; SDPs are indicated in blue.

In recent years, significant efforts have focused on challenging BB tasks. Several studies have been proposed that incorporate feature-based models. TEES [5], which adopted support vector machine (SVM) with a variety of features based on shortest dependency paths (SDPs), was the best-performing system with an F1 score of 42.27% in the BioNLP-ST’13 [4]. The VERSE team [6], which placed first in BioNLP-ST’16 with an F1 score of 55.80%, utilized SVM with rich features and a minimum spanning dependency tree (MST). Feature-based models, however, heavily depend on feature engineering, which sometimes is limited by its lack of domain-specific knowledge [7].

Since 2014, deep learning (DL) methods have garnered increasing attention due to their state-of-the-art performance in several natural language processing (NLP) tasks [8]. Unlike the feature-based models, DL models demand less feature engineering because they can automatically learn useful features from training data. Examples of popular DL models that have successfully been applied for biomedical relation extraction include Convolutional Neural Networks (CNNs) [9, 10, 11, 12] and Recurrent Neural Networks (RNNs) [13, 14].

Other than features-based models in the BB task, several previous studies using DL approaches have significantly outperformed traditional SVM approaches. For example, in BioNLP-ST’16, DUTIR [15] utilized CNN models to achieve an F1 score of 47.80%; TurkuNLP [16] used multiple long short-term memories (LSTM) with SDPs to achieve an F1 score of 52.10% and was ranked second in the competition. DET-BLSTM [17] applied bidirectional LSTM (BLSTM) with a dynamic extended tree (DET) adapted from SDPs and achieved an F1 score of 57.14%. Recently, BGRU-Attn [18] proposed bidirectional gated recurrent unit (BGRU) with attention mechanism and domain-oriented distributed word representation. Consequently, it became the state-of-the-art DL system without hand-designed features for the BB task with an F1 score of 57.42%.

Despite the success of DL in previous studies, there are still several limitations to be considered. Although SDPs have been shown to contain valuable syntactic features for relation extraction [16, 17, 18, 19, 20, 21], they still may miss some important information. For example, in Figure 1, the word “in”, which should play a key role in defining the relation between the bacteria “mycobacteria” and the biotope “Queensland” is not included in SDP (represented by blue lines) because there is no dependency path between “in” and any entities. To overcome the limitation of SDPs, some studies have used sequences of full sentences to extract biomedical relations from texts [22, 23, 24]. However, it is very difficult for DL models to learn enough features from only sequences of sentences. Instead of learning from full sentences, attention networks have demonstrated success in a wide range of NLP tasks [25, 26, 27, 28, 29, 30, 31]. In addition, BGRU-Attn [18] first used the Additive attention mechanism [29] for the BB task to focus on only sections of the output from RNN instead of the entire outputs and achieved state-of-the-art performance. Other attention techniques such as Entity-Oriented attention [30] and Multi-Head attention [31] still have not been explored for this task. From the aspect of word representation, traditional word embeddings [32, 33] only allow for single context-independent representation. This situation can lead to word sense ambiguation across various linguistic contexts [34]. Contextual representations of words [35] and sentences [36] based on language-understanding models addressed this problem and achieved state-of-the-art performance on general-purpose domain NLP tasks [35, 36, 37, 38, 39]. Nevertheless, [40] has shown that the word-embedding models pre-trained on a general-purpose corpus such as Wikipedia are not suitable for biomedical-domain tasks. Finally, the training process of DL approaches with many randomly initialized parameters is non-deterministic—multiple executions of the same model may not result in the same outcome. To solve this issue and provide a statistical comparison of models’ performances, [41, 42] reported the mean F1 score of the same model architecture initialized with different parameter settings (random seeds). This evaluation metric indicates the average behavior of a model’s performance and is more suitable for the biases and trends in real-world applications [43]. However, the mean F1 score had never been explored in prior studies of the BB task.

In this study, we propose a hybrid model between a RNN and a feed-forward neural network such as a CNN. We use the RNN to extract full-sentence features from long and complicated sentences. We also apply the CNN to capture SDP features that are shorter, more valuable, and more concise. In addition, because attention mechanisms have been proven to be helpful in the BB task [18], we incorporate several kinds of attention mechanisms—Additive attention, Entity-Oriented attention, and Multi-Head attention—into the model. Furthermore, we integrate domain-specific contextual word representation into the model to provide word-sense disambiguation. Sentence representation was also introduced to improve the full-sentence model by embedding sequence sentence information from a pre-trained language understanding model. To address the uncertainty of a single run model’s performance measured by the maximum F1 score, we used the mean F1 score as an evaluation metric for comparisons of the models.

## 2 Methods

In this section, we describe the proposed DL model for extracting BB relations from the given biomedical literature (Figure 2).

**Figure 2:**
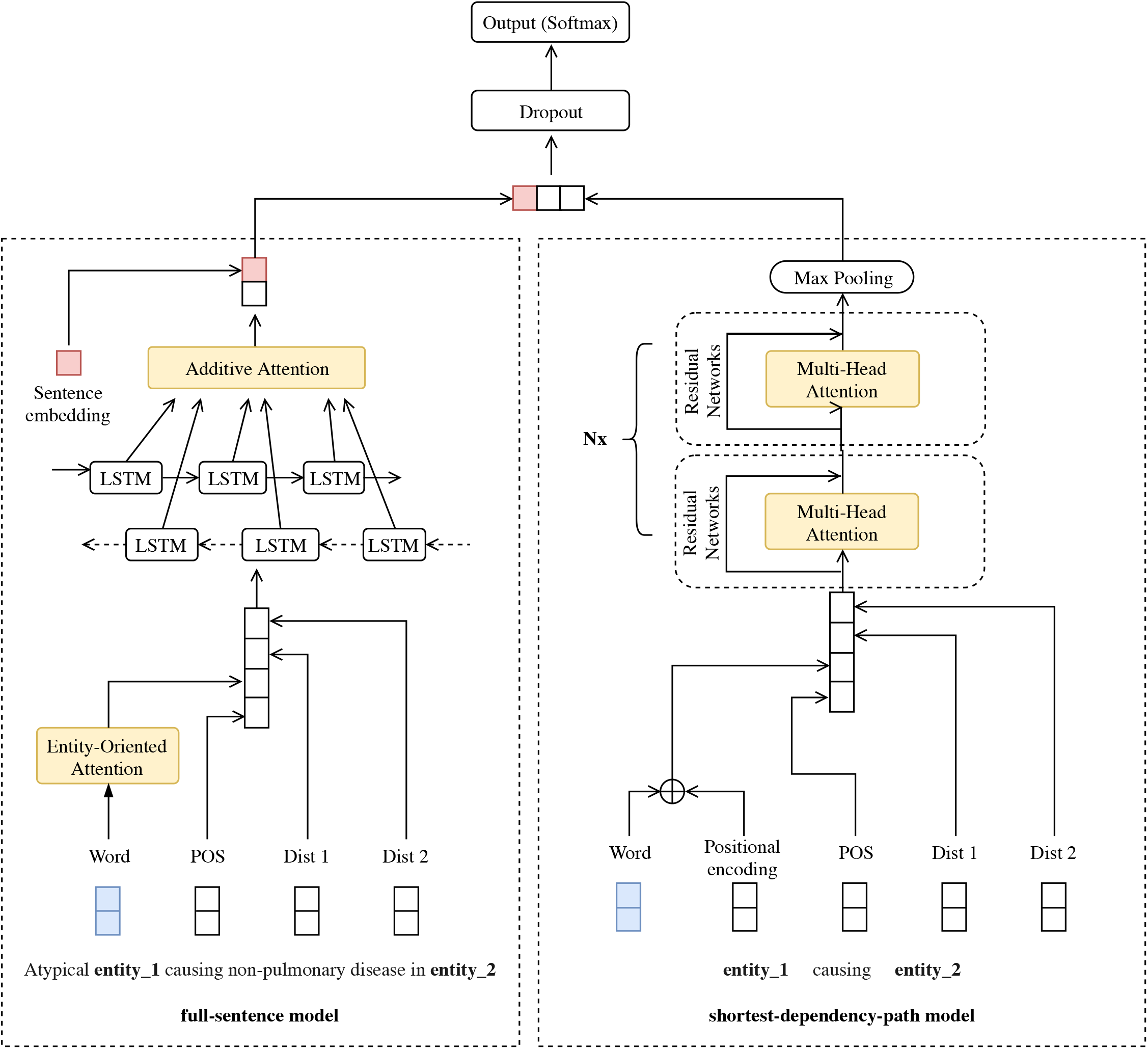
The overall architecture of our proposed model with the combined full-sentence and SDP models, together with various attention mechanisms.

### 2.1 Text preprocessing

We used the TEES system [5, 16] to run the pipeline of the text preprocessing steps. Tokenization and part-of-speech (POS) tagging for each word in a sentence were generated using the BLLIP parser [44] with the biomedical-domain model. The dependency grammar resulted from the BLLIP was further processed using the Stanford conversion tool [45] to obtain the Stanford dependencies (SD) graph.

We then used Dijkstra’s algorithm to determine the SDPs between each pair of entities: bacteria and biotope. The SDPs represented the most relevant information and diminished noises by undirected graph (Figure 1). An entity pair was neglected if there was no SDP between the entities. While the dependency paths only connect a single word to others within the same sentence (intra-sentence), there are some cross-sentence (inter-sentence) associations that can be very challenging in terms of the extraction task. In order to compare with other existing works [5, 15, 16, 17, 18], only intra-sentence relations were considered.

To ensure the generalization of the models, we followed the protocol of previous studies [17, 18] that blinded the entities in a sentence. The bacteria and location mentions were replaced by “entity_1” and “entity_2” respectively. For example, as shown in Table 1, we can generate two BB relation candidates (termed “instances”) from a sentence “Long-term ***Helicobacter pylori*** infection and the development of atrophic *gastritis* and gastric cancer in *Japan*.”, where the bacteria and location mentions are highlighted in bold italics and italics, respectively. After entity blinding, we converted all words to lowercase to simplify the searching process and improve text matching.

**Table 1:**
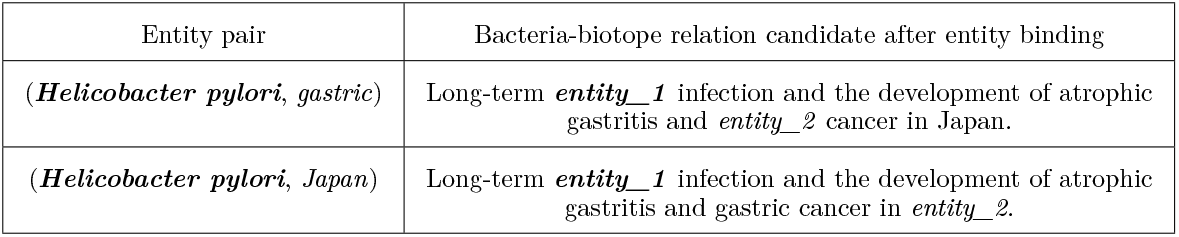
Bacteria-biotope relation candidates (instances) in a sentence after entity blinding. The bacteria and location mentions are highlighted in bold italics and italics, respectively.

### 2.2 Input embedding representations

The input representations used in our model were divided into full-sentence and SDP features. Let {*w*_1_, *w*_2_,…, *w_m_*} and {*s*_1_, *s*_2_,…, *s_n_*} denote the full sentence and SDPs of a sentence that are represented by different embeddings. Each word *w_i_* in a full sentence was represented by word vector, POS, and distance embeddings. Each word *s_j_* in the SDP was represented by word vector, POS, and distance embeddings together with positional encoding (PE). The detailed embeddings used in our model are explained below.

For a full sentence in the RNN model, **word embedding** was a 200-dimensional word vector, the pre-trained biomedical word-embedding model [46], built from a combination of PubMed and PMC texts using Word2Vec [32]. **Part-of-speech embedding** was initialized randomly at the beginning of the training phase.

**Distance embedding** [18, 47] is derived from the relative distances of the current word to the bacteria and location mentions. For example, in Figure 1, the relative distances of the word “in” to bacteria “mycobacteria” and location “Queensland” are −4 and 1, respectively. To construct the distance embedding *D*(*l*) for each relative distance, every dimension *d*(*l*) of the distance embedding is initialized using equation (1), where *l* is the relative distance and *s* refers to the maximum of the relative distances in the dataset. All *d*(*l*) dimensions form the distance vectors [*dist*_1_, *dist*_2_], which represent the distance embeddings *D*(*l*) of the current word to the bacteria and location mentions, respectively.

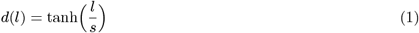

For SDP in the CNN model, we used **PE** [31] to inject some information about the absolute position of the words in the sentence. The PE vectors were initialized by sine and cosine functions of different frequencies; these functions embed information based on their relative position. Because PE has the same dimension as the word embedding, we can sum these two vectors.

In summary, the overall input embedding representation for a word *w_i_* in full sentences is 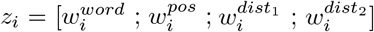. Similarly, for a given word *s_j_* on the SDP the overall input embedding representation is 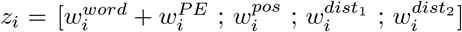.

### 2.3 A DL model based on full sentences and SDPs

#### 2.3.1 Full-sentence model

We employed BLSTM [48] to learn global features from full sentences. The BLSTM can be used to encode the sequential inputs both forward and backward, and it has been shown to outperform one-way LSTM in many studies [13, 47, 48, 49, 50]. Given a full sentence of *M* tokens, {*z*_1_, *z*_2_,…, *z_M_*}, at the *t*-th time step, the BLSTM takes the current input representation (*z_i_*), previous hidden state (*h*_*t*−1_), and previous memory cell (*c*_*t*−1_) as its inputs to generate the current hidden state (*h_i_*) and memory cell (*c_i_*).

#### 2.3.2 SDP model

The multiple-filter-widths CNN model [51] was proposed for SDP model to learn local features from SDPs. For a given SDP sequence of *N* tokens, {*z*_1_, *z*_2_,…, *z_N_*}, let *z_i_* ∈ ℜ^*k*^ be the *k*-dimensional input embedding vector corresponding to the *i*-th word in the sequence. The CNN takes an input sequence of length *N* to generate the feature map (*c_i_*) by convolutional filters and max pooling operations. Compared with LSTM, the CNN model is expected to be better at extracting high-quality features from short and concise SDPs [52].

### 2.4 Attention mechanisms

Attention mechanisms have recently been successfully applied to biomedical relation extraction tasks [14, 18, 30]. These attention networks are able to learn a vector of important weights for each word in a sentence to reflect its impact on the final result. We propose the use of three attention mechanisms—Additive, Entity-Oriented, and Multi-Head—to improve our relation extraction. Figure 2 shows how these attention mechanisms are integrated into our proposed DL model.

#### 2.4.1 Additive attention

The Additive attention focuses on sentence-level information. It was first used by [29] to improve neural machine translation and recently applied to the BB task [18]. The idea of Additive attention is to consider all LSTM hidden states with different attention weights when deriving the context vector. The context vector depends on the sequence of hidden states {*h*_1_, *h*_2_,…, *h_k_*}. Each hidden state contains information about the whole input sequence with a strong focus on the parts surrounding the *i*-th word. The context vector (*c_i_*) was computed as a weighted sum of these hidden states (*h_i_*) using equation (2). The attention weight (*a_i_*) of each hidden state (*h_j_*) was then computed using equation (3). The additive attention assigned a score (*a_i_*) to the pair of input at position *i*, which was parameterized using a feed-forward network with a single hidden layer. The model was then jointly trained with other parts of the model. The attention score function is shown in equation (4), where *v_a_* is the weight matrix to be learned.

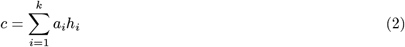

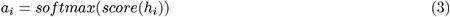

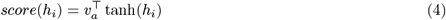

#### 2.4.2 Entity-Oriented attention

This attention focuses on word-level information. It has been used by [29, 31] and has been shown to be effective at drug-drug interaction (DDI) extraction [30]. This attention was used to determine which words in a sentence most influence the relationship between a pair of entities. This attention mechanism was applied after our word-embedding layer to quantify the concentration of word-level information. We exploited two attention weights (*a_j_*), *j* ∈ {1, 2}, which denoted the relevance degree of each word (*w_i_*) of a sentence with respect to the *j*-th entity mention (*e_j_*). The attention score 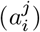 was calculated using the dot product operation (·) of the current word embedding vector (*u_w_i__*) and the *j*-th entity word-embedding vector (*u_e_j__*). The score was then normalized by the dimensionality of word embedding vector (*m*) using equation (5). The attention weight (*a_i_*) of word (*w_i_*) to both entities was computed using equation (6).

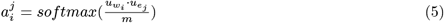

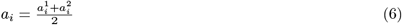

#### 2.4.3 Multi-Head attention

The Multi-Head attention focuses on extracting local features in multiple subspaces. It was used by [31] for machine translation. These authors showed that the model jointly attended to information from different representation subspaces at different positions. Instead of performing a single scale dot-product attention, [31] found to be beneficial if the queries (*q*), keys (*k*), and values (*v*) of the same dimension (*d_k_*) were linearly projected *h* times with different learned projections. For each head, we computed the dot products (·) of the queries of length (*n*) with all keys. We then divided each by 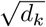 and applied the softmax function to obtain the attention score using equation (7). The context weight (*c_i_*) on the values was computed using equation (8). To obtain Multi-Head feature representations, the context weights from *h* heads were concatenated using equation (9). Compared with CNNs, Multi-Head feature representation (*h_out_*) uses *h* parallel attention mechanisms with different low-dimensional projection instead of a fixed-width convolutional filter. Inspired by the transformer network [31], stacks of Multi-Head attentions were employed in our model with residual connections and PE. Figure 3 shows the overview architecture of the CNN model and Multi-Head attentions as our SDP model.

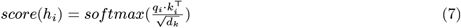

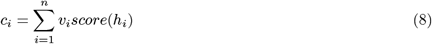

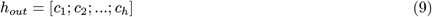

**Figure 3:**
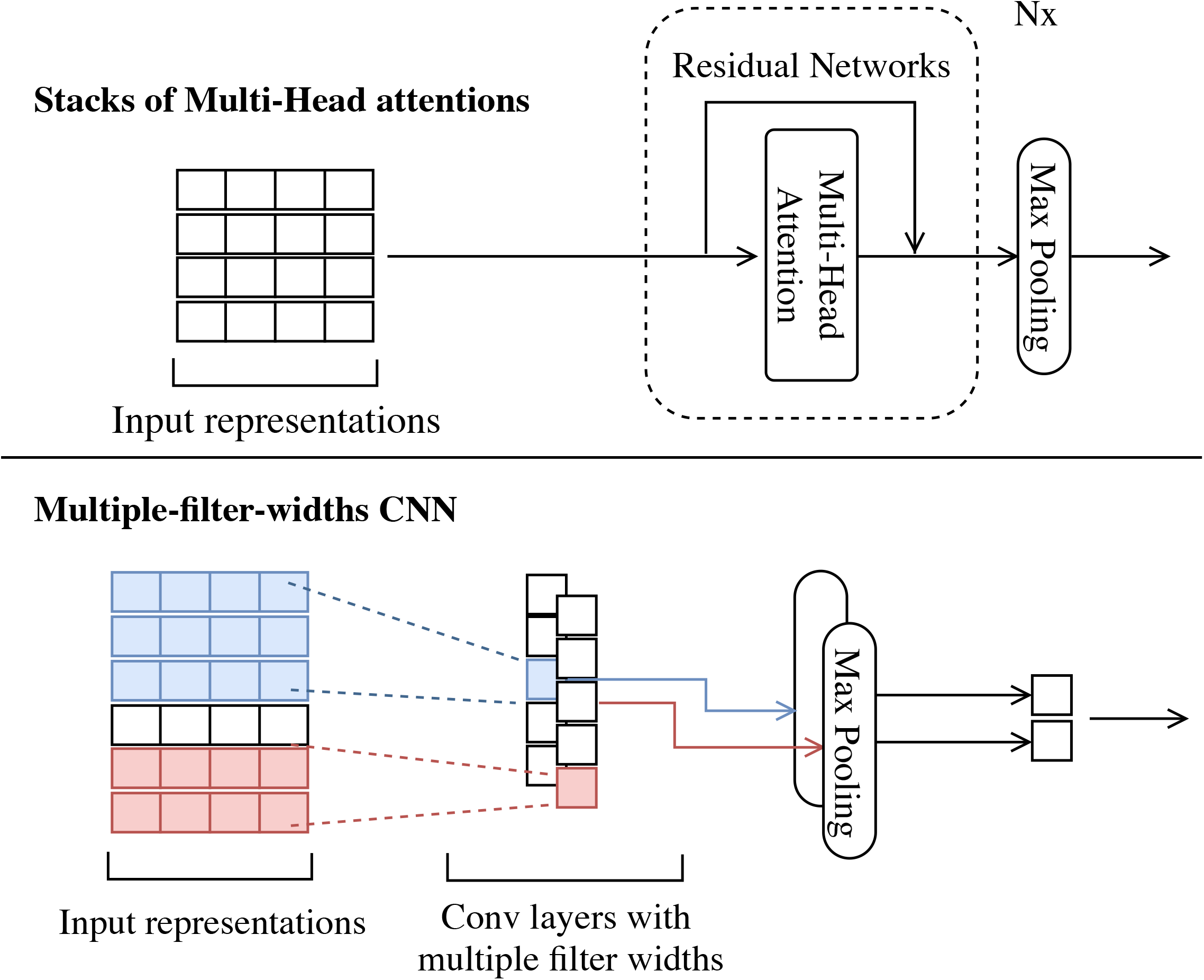
Illustration of the SDP model architecture to extract local features based on a CNN model or N stacks of Multi-Head attentions.

### 2.5 Contextual representations

The choice of how to represent words or sentences poses a fundamental challenge for NLP communities. There have been some advances in universal pre-trained contextual representations on a large corpus that can be plugged into a variety of NLP tasks to automatically improve their performance [35, 36]. By incorporating some contextualized information, these representations have been shown in [35, 36, 37, 38, 39] to alleviate the problem of ambiguation and outperform traditional context-free models [32, 33]. In this study, we propose two contextual embedding models pre-trained on a biomedical corpus of words and sentences.

#### 2.5.1 Contextual word representation

The contextual word vector used in our proposed model was generated by ELMo [35]. ELMo learned word representations from the internal states of a bidirectional language model. It was shown to improve the state-of-the-art models for several challenging NLP tasks. Context-free models such as Skip-gram [32] and GloVe [33] generate a single word representation for each word in their vocabulary. For instance, the word “cold” would have the same representation in “common cold” and “cold sensation” [34]. On the other hand, contextual models will generate a representation of the word “cold” differently based on context. This representation can be easily added to our proposed model by reconstituting the 200-dimensional word vectors with the new pre-trained contextual word vectors. Currently, the ELMo model, pre-trained on a large general-purpose corpus (5.5 billion tokens), is freely available to use [35]. However, [40, 53] showed that domain-irrelevant word-embedding models pre-trained on large, general-purpose collections of texts are not sufficient for biomedical-domain tasks. Therefore, we present a domain-specific, contextual, word-embedding model pre-trained on a bacterial-relevant corpus. Inspired by the relevance-based word embedding [54], the corpus to pre-train our proposed contextual word embedding model included relevance-based abstracts downloaded from PubMed, which contain only sentences with bacterial scientific names from the BB task (118 million tokens). To evaluate the effectiveness of our proposed domain-specific, contextual, word-embedding model, we compared it with the contextual model pre-trained on randomly selected abstracts from PubMed with the same number of tokens. All of the pre-trained models were fine-tuned with the BB dataset in order to transfer learned features from the pre-train models to our task.

#### 2.5.2 Contextual sentence representation

Our contextual sentence embedding was constructed by BERT [36]. BERT represents words based on a bidirectional approach and learns relationships between sentences. Hence, BERT representation unambiguously represents both words and sentences. However, due to the limited computational resource to pre-train BERT using our biomedical corpus the available pre-trained BERT on general-purpose corpus was adopted and fine-tuned with the BB task.

### 2.6 Training and classification

The output layer used the softmax function [55] to classify the relationship between pairs of bacteria and biotope mentions. The softmax layer takes the output of BLSTM for full-sentence feature, the output of Multi-Head attention networks for SDP feature, and the sentence embedding from BERT as its inputs (Figure 2). These inputs are fed into a fully connected neural network. The softmax layer’s output was the categorical probability distribution over each class type (*c*) by equation (10).

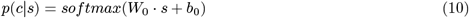

where *W*_0_ and *b*_0_ are weight parameters and *s* is the feature representation of sentences. For the binary classification, we used the cross-entropy cost function (*J*(*θ*)) as the training objective by equation (11).

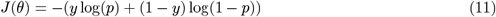

where *y* is the binary indicator (0 or 1) if the class label is correct for each predicted sentence and *p* is the predicted probability. Additionally, we applied Adam optimization to update the network weights with respect to the cost function.

### 2.7 Data

#### 2.7.1 Training and test datasets

The dataset provided by the BB task [2] of BioNLP-ST’16 consists of titles and abstracts from PubMed with respect to reference knowledge sources (NCBI taxonomy and OntoBiotope ontology). All entity mentions—*Bacteria, Habitat*, and *Geographical*—and their interactions were manually annotated from diverse-backgrounds annotators. Each bacteria-biotope pair was annotated as either a negative or positive *Lives_in* relation. The relations can be defined as inter-sentence and intra-sentence. In our study, we also followed previous studies [5, 15, 16, 17, 18] in simply excluding inter-sentence instances from the dataset. This procedure resulted in the removal of 107 and 64 annotated instances from the training data and development data, respectively. Table 2 lists the statistics of the preprocessed BB dataset used in our experiments.

**Table 2:** Statistics of a preprocessed BB dataset.

|  | Train | Dev | Test |
| --- | --- | --- | --- |
| Avg.word in SDPs | 4.81 | 4.86 | 4.58 |
| Avg.word in full sentences | 27.22 | 29.92 | 28.12 |
| Positive instances | 248 | 173 | - |
| Negative instances | 275 | 332 | - |
| Total instances | 523 | 505 | 532 |

#### 2.7.2 The pre-training corpus of contextual word representations

In order to get the proposed domain-specific word embeddings (specific-PubMed ELMo), we pre-trained ELMo on the bacterial-relevant abstracts downloaded from the PubMed database. These specific abstracts contain roughly 118 million words that use all of the bacteria names that are noted in the BB dataset as keywords. An example keyword is the bacteria mention “mycobacteria” (Figure 1). Furthermore, we pre-trained another domain-general word embeddings (random-PubMed ELMo) on randomly selected PubMed abstracts with a similar corpus size to evaluate the performance of the domain-specific model. To reduce the memory requirement of both pre-training models, we only used the words in the training, development, and test sets to construct the vocabularies.

### 2.8 Hyper-parameter setting

We used the Pytorch library [56] to implement the model and empirically tuned the hyper parameters using 3-fold cross validation on the training and development data. After tuning, the dimensions of the contextual word embedding (ELMo), context-free word embedding, POS embedding, distance embedding, and sentence embedding (BERT) were 400, 200, 100, 300, and 768, respectively. The dimension of PE was set to either 200 or 400 for the context-free or contextual word embeddings, respectively. The hidden unit numbers of BLSTM and the filter numbers of CNN were 64. The convolutional window sizes were 3, 5, and 7. For the Multi-Head attention mechanism, we used three stacks of Multi-Head attentions with respect to the residual connections; the number of heads for each stack was 2. Before the output layer, we applied a dropout rate of 0.5 to the concatenation of full-sentence, SDP, and sentence-embedding features. The mini-batch was set to 4, and a rectified linear unit (ReLU) was used as our activation functions. We set the learning rate to 0.001 for Adam optimization. Despite the underfitting problem, we used different parameters of the only full-sentences model, denoted as “full-sentence” in the Influence of full-sentence and SDP features section. The dropout rate was set to 0.1, and the hidden unit number of LSTM was 32.

### 2.9 Evaluation metrics

For our model, the final results on the test dataset were evaluated using the online evaluation service provided by the BB task of the BioNLP-ST’16. Due to the removal of inter-sentence examples, any inter-sentence relations in the test dataset that counted against our submission were considered to be false negatives.

As discussed above, a large number of local minima in DL models can lead to larger parameter spaces. Evaluating a single model several times tends to result in performance convergence under different parameter initializations (or random seeds). To alleviate this problem, we reported the mean F1 score instead of only the maximum F1 score reported by previous studies [5, 6, 15, 16, 17, 18]. To calculate the mean F1 score, we built 30 models as suggested by [41]. These models were trained using the same architecture but with different random seeds. Then, we evaluated the F1 score of each model on the same test set using an online evaluation service. With these F1 scores, we then calculated the minimum, maximum, mean, and standard deviation to assess the robustness of the model. In this study, we used the mean F1 score as the main evaluation metric; the maximum F1 score was still used to compare with other previously used models.

## 3 Results

We assessed the performance of our model as follows. First, we compared our model with existing models in terms of maximum and average F1 scores. Then, we evaluated the effectiveness of each contribution used by the model: feature combination between full sentences and SDP, attention mechanisms, contextual word representation, and contextual sentence representation. Here, we discuss the overall experimental results of this proposed model.

### 3.1 Performace comparisons with existing models

#### 3.1.1 Maximum F1 score comparisons

Table 3 lists the maximum F1 score of our model compared with those of prior studies. In the BB task [2], each team evaluated the model on the test set using an online evaluation service. Most of the existing systems were based either on SVM or DL models. The SVM-based baseline [5] was a pipeline framework using SVMs on SDPs with an F1 score of 42.27%. Similarly, [6] proposed a utilized SVM with rich feature selection that yielded an F1 score of 55.80%. Compared with SVM-based models, DL-based models automatically learn feature representations from sentences and achieve state-of-the-art performance. For example, DUTIR [15] utilized a multiple-filter-widths CNN to achieve an F1 score of 47.80%. TurkuNLP [16] employed a combination of several LSTMs on the shortest dependency graphs to obtain the highest precision of 62.30% and an F1 score of 52.10%. BGRU-Attn [18] proposed a bidirectional GRU with the attention mechanism and biomedical-domain-oriented word embedding to achieve the highest recall of 69.82% and an F1 score of 57.42%. These results reveal that our proposed model achieved the best performance in the official evaluation (i.e., the highest F1 score: 60.77%). In contrast with the previous state-of-the-art model (BGRU-Attn [18]), our model achieved more balanced precision (56.85%) and recall (65.28%). The results revealed that our model could leverage both full-sentence and SDP models along with contextual representations to capture the vital lexical and syntactic features of given sentences. Therefore, our model can combine the advantages of all contributions to achieve a good trade-off between precision and recall, which resulted in its superior performance in the BB corpus.

**Table 3:** Performance comparison (maximum F1 score) with existing models. The listed results derive from the corresponding papers. F: F1 score; R: recall; P: precision. Our model used all of the proposed contributions (the results from the last row in Table 8). The highest scores are highlighted in bold.

| Methods | Team | F | R | P |
| --- | --- | --- | --- | --- |
| SVM-based | TEES [5] | 42.27 | 38.35 | 61.61 |
|  | VERSE [6] | 55.80 | 61.50 | 51.00 |
| DL-based | DUTIR [15] | 47.80 | 39.70 | 60.00 |
|  | TurkuNLP [16] | 52.10 | 44.80 | **62.30** |
|  | DET-BLSTM [17] | 57.14 | 57.99 | 56.32 |
|  | BGRU-Attn [18] | 57.42 | **69.82** | 48.76 |
|  | **Our model** | **60.77** | 65.28 | 56.85 |

#### 3.1.2 Mean F1 score comparisons

Table 4 lists the results of our model compared with other DL models: TurkuNLP [16] and BGRU-Attn [18] based on mean F1 scores. Our model achieved the highest mean F1 score and the lowest standard deviation. This finding indicates that our model is more robust to randomness and highly consistent in its performance. To provide a statistically significant comparison of our model’s performance, we also performed a two-sample t-test with the hypothesis that two populations (our model and a compared model) were equal in terms of their mean F1 scores (null hypothesis *H*_0_). The results revealed that we rejected the null hypothesis with a p-value less than 0.001 (or more than 99.9% confidence). This fact implied that our model’s mean F1 score was significantly better than that of other models.

**Table 4:** Performance comparison (mean F1 score) with existing models. These results, from existing models, derive from the model reimplementation. The highest scores are highlighted in bold except for the standard deviation (SD). The p-value was calculated using the two-sample t-test for the difference of mean F1 score.

| Model | F1 score |  |  |  | p-value |
| --- | --- | --- | --- | --- | --- |
|  | Mean | SD | Min | Max |  |
| TurkuNLP [16] | 46.18 | 4.64 | 35.07 | 51.99 | <0.001 |
| BGRU-Attn [18] | 50.22 | 2.79 | 43.05 | 55.54 | <0.001 |
| **Our model** | **57.63** | **1.15** | **54.41** | **60.77** | - |

### 3.2 Effects analysis of each proposed strategy

In the following sections, we evaluate the effectiveness of each model contribution: combined full-sentence and SDP models, attention mechanisms, contextual word representation, and contextual sentence representation (Tables 5–8). To overcome the variant problem in model evaluation, each experiment used the mean F1 score for model selection and evaluation.

**Table 5:** The effectiveness of the application of full-sentence and SDP features according to the mean F1 scores of 30 different random seeds. All of the highest scores are highlighted in bold except for the standard deviation (SD).

| Model |  | F1 score |  |  |  |
| --- | --- | --- | --- | --- | --- |
| Fulls | SDPs | Mean | SD | Min | Max |
| - | CNN | 43.79 | 3.39 | 37.02 | 51.82 |
| BLSTM | - | 41.22 | 14.49 | 12.82 | 49.93 |
| **BLSTM** | **CNN** | **45.96** | **2.87** | **42.09** | **52.19** |

**Table 6:**
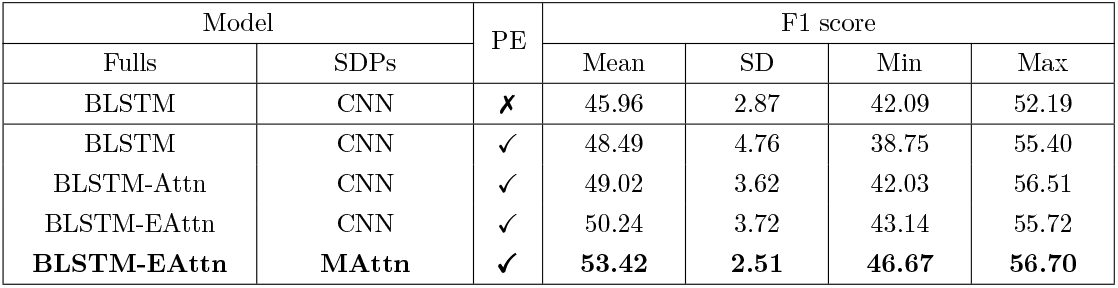
The effectiveness of the integrated attention mechanisms according to mean F1 scores for 30 different random seeds. All of the highest scores are highlighted in bold except for the standard deviation (SD). The first-row results derive from the best results of previous experiments (i.e., the last row in Table 5). Note: “PE” denotes positional encoding, “Attn” denotes the use of only Additive attention, “EAttn” denotes the use of both Additive and Entity-Oriented attentions, and “MAttn” denotes the use of Multi-Head attention.

| Model |  | PE | F1 score |  |  |  |
| --- | --- | --- | --- | --- | --- | --- |
| Fulls | SDPs |  | Mean | SD | Min | Max |
| BLSTM | CNN | **✗** | 45.96 | 2.87 | 42.09 | 52.19 |
| BLSTM | CNN | ✓ | 48.49 | 4.76 | 38.75 | 55.40 |
| BLSTM-Attn | CNN | ✓ | 49.02 | 3.62 | 42.03 | 56.51 |
| BLSTM-EAttn | CNN | ✓ | 50.24 | 3.72 | 43.14 | 55.72 |
| **BLSTM-EAttn** | **MAttn** | **✓** | **53.42** | **2.51** | **46.67** | **56.70** |

**Table 7:** The effectiveness of domain-specific contextual word representation according to the mean F1 scores of 30 different random seeds. All of the highest scores are highlighted in bold except for the standard deviation (SD). The first-row results derive from the best results of previous experiments (i.e., the last row in Table 6). Note: “PubMed word2vec” denotes the context-free word model, “general-purpose ELMo” denotes the general-purpose contextual word model, “random-PubMed ELMo” denotes the domain-general contextual word model based on 118 million randomly selected tokens abstracts from PubMed, and “specific-PubMed ELMo” denotes the domain-specific contextual word model based on 118 million bacterial-relevant abstracts from PubMed.

| Pre-trained word model | F1 score |  |  |  |
| --- | --- | --- | --- | --- |
|  | Mean | SD | Min | Max |
| PubMed word2vec | 53.42 | 2.51 | 46.67 | 56.70 |
| general-purpose ELMo | 54.30 | 3.61 | 42.76 | 56.51 |
| random-PubMed ELMo | 53.81 | 3.65 | 38.89 | 57.01 |
| **specific-PubMed ELMo** | **55.91** | **1.49** | **51.24** | **57.48** |

**Table 8:** The effectiveness of the contextual sentence representation by the mean F1 scores of 30 different random seeds. All of the highest scores are highlighted in bold except for the standard deviation (SD). The first-row results derive from the best results of previous experiments (i.e., the last row in Table 7).

| Sentence representation | F1 score |  |  |  |
| --- | --- | --- | --- | --- |
|  | Mean | SD | Min | Max |
| without | 55.91 | 1.49 | 51.24 | 57.48 |
| **with** | **57.63** | **1.15** | **54.41** | **60.77** |

#### 3.2.1 Influence of full-sentence and SDP features

Table 5 lists the mean F1 score of 30 DL models with different random seeds. The mean F1 score obtained from the experiment indicated that the use of full-sentence and SDP models together outperformed the separated models. The data in Table 5 also demonstrate that CNN achieved better performances than BLSTM when BLSTM and CNN were separately applied to the full sentences and SDPs, respectively. This result suggests that our model effectively combines the SDP and full-sentence models to extract more valuable lexical and syntactic features. These features were generated not only from two different sequences (full sentences and SDPs) but also two different neural network structures (BLSTM and CNN).

#### 3.2.2 Influence of Attention mechanisms

After we measured the effectiveness of the full-sentence and SDP features, we additionally explored the effects of the Additive, Entity-Oriented, and Multi-Head attention mechanisms. The attention mechanisms were applied to concentrate the most relevant input representation instead of focusing on entire sentences. Table 6 lists the productiveness of each attention mechanism integrated into our full-sentence and SDP models. According to [31], Multi-Head attention networks were first proposed with the use of PE to insert valuable locality information. Because Multi-Head attention networks were employed with PE, we applied PE to CNN in order to fairly compare the effectiveness of Multi-Head attention. The use of the Additive attention mechanism improved the mean F1 score by 0.53%. Entity-Oriented attention improved the average F1 score from 49.02 to 50.24%. These results show that attention mechanisms might highlight influential words for the annotated relations and help reveal semantic relationships between each entity. This approach improved the overall performance of our model. Finally, the stacks of Multi-Head attention networks were the primary contributor to our model. The experimental results revealed that the proposed model using Multi-Head attention together with SDPs increased the mean F1 score by 3.18% compared with the proposed model using CNN. Our proposed model used stacks of Multi-Head attentions with residual connections instead of CNN, as shown in the overall architecture of Figure 2.

#### 3.2.3 Influence of domain-specific contextual word representation

Table 7 lists the effectiveness of our domain-specific, contextual word representation to our model after previous contributions (combined features and attention mechanisms). The contextual word representation (ELMo) was proposed to provide word sense disambiguation across various linguistic contexts and handle out-of-vocabulary (OOV) words using a character-based approach. The results in Table 7 reveal that every ELMo model outperformed the traditional word2vec model. One possible explanation for this finding is that the ELMo model uses a character-based method to handle OOV words while word2vec initializes these OOV word representations randomly. The ELMo model can also efficiently encode different types of syntactic and semantic information about words in context and therefore improve overall performance. The use of our proposed contextual word model with a domain-specific corpus (specific-PubMed ELMo) achieved the highest average F1 score of 55.91%. This score represented an improvement by 2.49%, 1.61%, and 2.10% compared with the score deriving from the use of PubMed word2vec, general-purpose ELMo, and random-PubMed ELMo, respectively. These improvements reveal the importance of taking relevant information into account when training contextual embedding vectors. We also noted that the general-purpose ELMo achieved slightly better performance compared with the random-PubMed ELMo. However, the latter was pre-trained on a biomedical-domain corpus; the size of the pre-trained corpus of the former (5.5 billion tokens) is significantly larger than that of the latter (118 million tokens), which resulted in the higher-quality word embeddings and better semantic representations.

#### 3.2.4 Influence of contextual sentence representation

In order to use sentence embeddings as fixed features from the pre-trained BERT, [36] suggested that the best-performing method involved concatenating the feature representations from the top four 768-dimensional BLSTM hidden layers of the pre-trained model. However, we found that it was better to sum up the last four 768-dimensional hidden layers into the 768-dimension sentence embedding. This situation may have been due to the small training dataset. The addition of contextual sentence representation from the fine-tuned BERT model improved the mean F1 score by 1.68% (Table 8). The results suggest that the fine-tuned BERT model could enhance the full-sentence model to encode crucial contextual representations of long and complicated sentences.

### 3.3 Discussion

Our proposed model can take advantage of the proposed contributions in order to construct rich syntactic and semantic feature representations. Our model significantly outperforms other existing models in terms of both mean F1 score (57.63%; SD = 1.15%) and maximum F1 score (60.77%). The mechanisms that largely support stable performance include the Multi-Head attentions and domain-specific contextual word representation, which are responsible for mean F1 score increases of 3.18% and 2.49%, respectively. A possible advantage of Multi-Head attention compared with CNN is the ability to determine the most relevant local feature representations from multiple subspaces to the BB task based on attention weights. In addition, domain-specific contextual word representation is beneficial to the proposed model for capturing contextual embeddings from a bacterial-relevant corpus. The box-and-whisker plot in Figure 4 shows the mean F1 score distribution of the existing DL models and our final proposed model (blue boxes). The boxplot illustrates the performance of our model after incrementally adding each of the main contributions (grey boxes). The mean F1 score of each model is shown as a line. The blue boxes indicate the comparison of our final model and two reimplemented TurkuNLP [16] and BGRU-Attn [18]. The mean F1 score of our model was 57.63%, which exceeds that of the TurkuNLP and BGRU-Attn models by 11.45% and 7.41%, respectively. In other words, our proposed model generally achieves better performance in terms of both mean and maximum F1 scores. Furthermore, the inter-quartile range of our proposed model is much smaller than that of other DL models. This finding demonstrates that the performance of our model is more robust and suitable for real-world applications.

**Figure 4:**
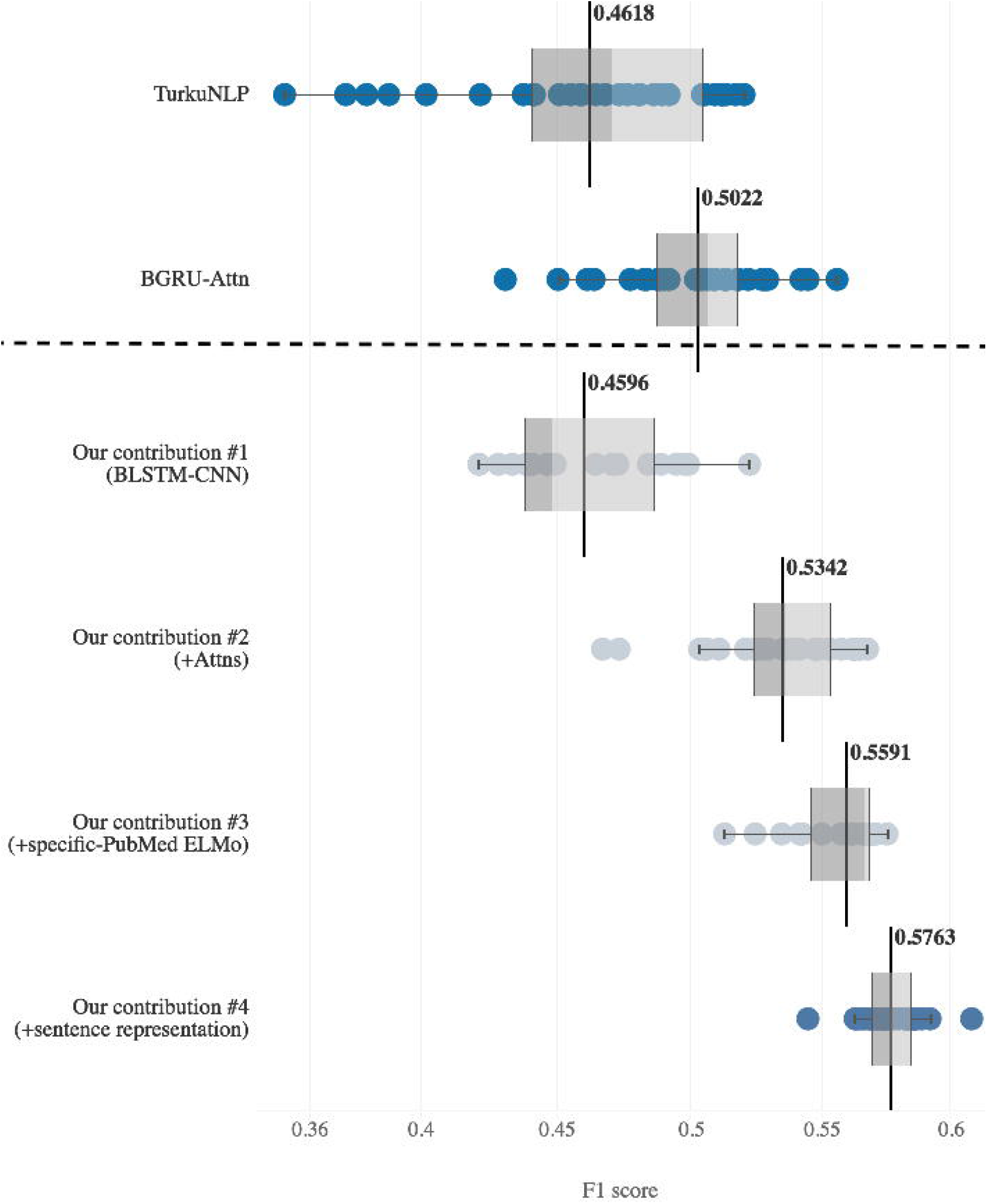
Box-and-whisker plot of average F1 score distributions of the deep-learning-based relation extraction models on the BB task. The comparison between our model and existing deep-learning-based models is shown in blue; the improvement of our model after adding each of the proposed contributions is shown in grey. Note: “Attns” denotes the use of integrated attention mechanisms.

For binary classification problems, F1 score is a common metric for evaluating an overall model’s performance because it conveys both precision and recall into one coherent metric. In some applications, however, it is more important to correctly classify instances than to obtain highly convergent results (i.e., high precision). On the other hand, some other applications place more emphasis on convergence rather than correctness (high recall). We experimented with using a frequency cut-off to explore how the probabilities output by the model function as a trade-off between precision and recall. Figure 5 shows the precision-recall curve (PRC) of our proposed model. When applied to real-world scenarios, users of the model are responsible for choosing the right cut-off value for their applications. For example, in semi-automated text-mining applications for knowledge management researchers never want to miss any bacteria-biotope relations. As a result, models with a high recall will be chosen to prescreen these relations. On the other hand, automated text-mining applications for decision support systems will require more precise relations. In Figure 5, our model with the default (0.5) cut-off value achieved an F1 score of 60.77% with balanced 56.85% recall and 65.28% precision. With a cut-off of 0.025, our model achieved the highest recall at 70.54% with 50.11% precision and an F1 score of 58.59%. With this cut-off value, our model outperformed the existing highest-recall model (BGRU-Attn [18]) by both 0.72% recall and 1.35% precision. Similarly, the line plot shown in Figure 5 shows that our model with a 0.975 cut-off achieved the highest precision (72.60%), recall (46.90%) and F1 score (56.99%). This model also outperformed the existing highest-precision model (TurkuNLP [16]) by 10.30% in precision and 2.10% in recall.

**Figure 5:**
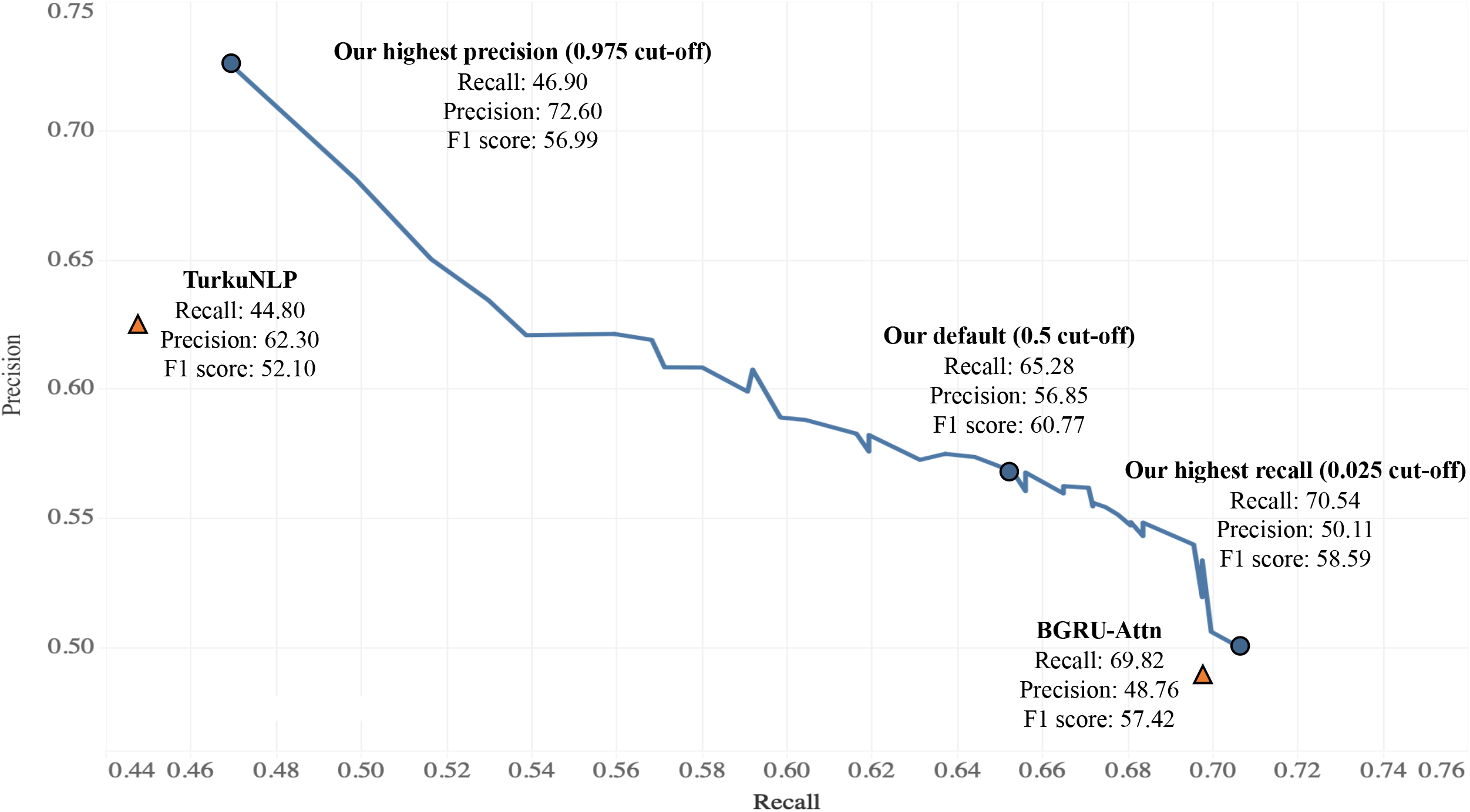
The precision-recall curve for our proposed model showing the trade-off between the true positive rate and the positive predictive value for our model using different probability thresholds (cut-off values).

To determine the factors that adversely affected the performance of our proposed model, we manually analyzed the correct and incorrect predictions from a development set compared with other existing models. We found that the proposed model could detect true negatives (TNs) better than other reimplemented models. This finding arose mainly because full-sentence features boosted the model’s ability to predict an entity pair as a false relation. For example, the sentence “*Rickettsia felis* was the only **entity_1** found infecting fleas, whereas *Rickettsia bellii* was the only agent infecting ticks, but no animal or human **entity_2** was **shown** to contain rickettsial DNA.”, where SDP are shown in bold, was predicted to be a false relation by our model. Other models predicted this sentence to be a true relation because of the word “shown” in the SDP. In addition, we found that false positives (FPs) were generally caused by the complex and coordinate structures of full sentences. A complicated sentence and a long distance between two entities can lead to relation classification failures. Examples of these adverse effects include the sentences “The 210 isolates with typical LPS patterns (119 Ara- clinical, 13 Ara- soil, 70 **entity_1 entity_2**, and 8 reference National Type Culture Collection strains) also exhibited similar immunoblot profiles against pooled sera from patients with melioidosis and hyperimmune mouse sera.” and “Testing animal and human sera by indirect immunofluorescence assay against four rickettsia antigens (*R. rickettsii, R. parkeri, R. felis*, and *R. bellii*), some opossum, **entity_2**, horse, and human sera reacted to **entity_1** with titers at least four-fold higher than to the other three rickettsial antigens.” In each of these sentences, the SDPs are highlighted in bold.

### 3.4 Limitations of our model

One of the most important limitations of our model is that it cannot extract inter-sentence relations between the bacteria and the biotopes. Hence, all true inter-sentence relations become false negatives. Inter-sentence relation extraction is much more challenging because it requires a more nuanced understanding of language to classify relations between entities in different sentences and clauses characterized by complex syntax. Because the size of our BB dataset is quite small, it is very difficult for DL models to learn sufficient high-quality features for the target tasks. However, this challenging task is left for future work. Furthermore, there is a large repertoire of biomedical literature and domain resources that are freely accessible and can be used as unlabeled data for semi-supervised learning and transfer learning methods [57, 58, 59].

## 4 Conclusions

We have presented a DL extraction model for the BB task based on a combination of full-sentence and SDP models that integrate various attention mechanisms. Furthermore, we introduced a pre-trained, contextual, word-embedding model based on the large bacteria-relevant corpus and fine-tuned contextual sentence representation. These embeddings encouraged the model to effectively learn high-quality feature representations from pre-trained language modeling. We evaluated our proposed model based on maximum and mean F1 scores. The experimental results demonstrated that our model effectively integrated these proposed contributions. The results showed that we could improve the performance of relation extraction to achieve the highest maximum and average F1 scores (60.77% and 57.63%, respectively). Our proposed model significantly outperformed other state-of-the-art models. Additionally, our model is more robust to real-world applications than the previous RE models.

Despite our model exhibiting the best performance on the BB task, some challenges remain. In particular, inter-sentence relations between bacteria and location entities have not been taken into account by any existing deep-learning-based models; this situation is likely due to insufficient training data. In the future, we plan to develop a new approach to increase the quantity and quality of limited training data for the target task using transfer learning and semi-supervised learning methods.

## References

[1] K. Bretonnel Cohen and Lawrence Hunter. Getting started in text mining. PLoS Computational Biology, 4(1):0001–0003, 2008.

[2] Louise Deléger, Robert Bossy, Estelle Chaix, Mouhamadou Ba, Arnaud Ferré, Philippe Bessì Eres, and Claire Nédellec. Overview of the Bacteria Biotope Task at BioNLP Shared Task 2016. Proceedings of the 4th BioNLP Shared Task Workshop, pages 12–22, 2016.

[3] Robert Bossy, Julien Jourde, Philippe Bessières, Maarten Van De Guchte, and Claire Nédellec. Bionlp shared task 2011: bacteria biotope. Proceedings of the BioNLP Workshop at ACL Conference, pages 56–64, 2011.

[4] Robert Bossy, Wiktoria Golik, Zorana Ratkovic, Philippe Bessières, and Claire Nédellec. BioNLP shared Task 2013–An Overview of the Bacteria Biotope Task. Proceedings of the BioNLP Workshop at ACL Conference, pages 153–160, 2013.

[5] Jari Björne and Tapio Salakoski. TEES 2.1: Automated Annotation Scheme Learning in the BioNLP 2013 Shared Task. BioNLP Shared Task 2013 Workshop, 2013:16–25, 2013.

[6] Jake Lever and Steven J. M. Jones. VERSE: Event and Relation Extraction in the BioNLP 2016 Shared Task. Proceedings of the 4th BioNLP Shared Task Workshop, pages 42–49, 2016.

[7] Yann LeCun, Yoshua Bengio, and Geoffrey Hinton. Deep learning. Nature, 521(7553):436–444, 5 2015.

[8] Shantanu Kumar. A Survey of Deep Learning Methods for Relation Extraction. 2017.

[9] Shengyu Liu, Buzhou Tang, Qingcai Chen, and Xiaolong Wang. Drug-Drug Interaction Extraction via Convolutional Neural Networks. Computational and Mathematical Methods in Medicine, 2016, 2016.

[10] Zhehuan Zhao, Zhihao Yang, Hongfei Lin, Jian Wang, and Song Gao. A protein-protein interaction extraction approach based on deep neural network. International Journal of Data Mining and Bioinformatics, 15(2):145, 2016.

[11] Daojian Zeng, Kang Liu, Siwei Lai, Guangyou Zhou, and Jun Zhao. Relation Classification via Convolutional Deep Neural Network. In Proceedings of the 25th International Conference on Computational Linguistics (COLING’14), 2014.

[12] Chanqin Quan, Lei Hua, Xiao Sun, and Wenjun Bai. Multichannel convolutional neural network for biological relation extraction. BioMed Research International, 2016.

[13] Sunil Kumar Sahu and Ashish Anand. Drug-drug interaction extraction from biomedical texts using long short-term memory network. Journal of Biomedical Informatics, 86(August):15–24, 2018.

[14] Yijia Zhang, Wei Zheng, Hongfei Lin, Jian Wang, Zhihao Yang, and Michel Dumontier. Drug-drug interaction extraction via hierarchical RNNs on sequence and shortest dependency paths. Bioinformatics, 34(5):828–835, 2018.

[15] Honglei Li, Jianhai Zhang, Jian Wang, Hongfei Lin, and Zhihao Yang. DUTIR in BioNLP-ST 2016: Utilizing Convolutional Network and Distributed Representation to Extract Complicate Relations. pages 93–100, 2016.

[16] Farrokh Mehryary, Jari Björne, Sampo Pyysalo, Tapio Salakoski, and Filip Ginter. Deep Learning with Minimal Training Data: TurkuNLP Entry in the BioNLP Shared Task 2016. Acl 2016, page 73, 2016.

[17] Lishuang Li, Jieqiong Zheng, Jia Wan, Degen Huang, and Xiaohui Lin. Biomedical event extraction via Long Short Term Memory networks along dynamic extended tree. In Proceedings - 2016 IEEE International Conference on Bioinformatics and Biomedicine, BIBM 2016, pages 739–742, 2017.

[18] Lishuang Li, Jia Wan, Jieqiong Zheng, and Jian Wang. Biomedical event extraction based on GRU integrating attention mechanism. BMC Bioinformatics, 2018.

[19] Yang Liu, Furu Wei, Sujian Li, Heng Ji, Ming Zhou, and Houfeng Wang. A Dependency-Based Neural Network for Relation Classification. (2006), 2015.

[20] Makoto Miwa and Mohit Bansal. End-to-End Relation Extraction using LSTMs on Sequences and Tree Structures. 2016.

[21] Xu Yan, Lili Mou, Ge Li, Yunchuan Chen, Hao Peng, and Zhi Jin. Classifying Relations via Long Short Term Memory Networks along Shortest Dependency Path. 2015.

[22] Suncong Zheng, Yuexing Hao, Dongyuan Lu, Hongyun Bao, Jiaming Xu, Hongwei Hao, and Bo Xu. Joint entity and relation extraction based on a hybrid neural network. Neurocomputing, 257:59–66, 2017.

[23] Ngoc Thang Vu, Heike Adel, Pankaj Gupta, and Hinrich Schütze. Combining Recurrent and Convolutional Neural Networks for Relation Classification. 2016.

[24] Yijia Zhang, Hongfei Lin, Zhihao Yang, Jian Wang, Shaowu Zhang, Yuanyuan Sun, and Liang Yang. A hybrid model based on neural networks for biomedical relation extraction. Journal of Biomedical Informatics, 81(March):83–92, 2018.

[25] Peng Zhou, Wei Shi, Jun Tian, Zhenyu Qi, Bingchen Li, Hongwei Hao, and Bo Xu. Attention-Based Bidirectional Long Short-Term Memory Networks for Relation Classification. In Proceedings of the 54th Annual Meeting of the Association for Computational Linguistics (Volume 2: Short Papers), 2016.

[26] Yong Zhang, Meng Joo Er, Rajasekar Venkatesan, Ning Wang, and Mahardhika Pratama. Sentiment classification using Comprehensive Attention Recurrent models. In Proceedings of the International Joint Conference on Neural Networks, 2016.

[27] Zhiwei Zhao and Youzheng Wu. Attention-based convolutional neural networks for sentence classification. In Proceedings of the Annual Conference of the International Speech Communication Association, INTERSPEECH, 2016.

[28] Yunfei Long, Lu Qin, Rong Xiang, Minglei Li, and Chu-Ren Huang. A Cognition Based Attention Model for Sentiment Analysis. In Proceedings of the 2017 Conference on Empirical Methods in Natural Language Processing, 2017.

[29] Minh-Thang Luong, Hieu Pham, and Christopher D Manning. Effective Approaches to Attention-based Neural Machine Translation. 2015.

[30] Wei Zheng, Hongfei Lin, Ling Luo, Zhehuan Zhao, Zhengguang Li, Yijia Zhang, Zhihao Yang, and Jian Wang. An attention-based effective neural model for drug-drug interactions extraction. BMC Bioinformatics, 18(1):1–11, 2017.

[31] Ashish Vaswani, Noam Shazeer, Niki Parmar, Jakob Uszkoreit, Llion Jones, Aidan N. Gomez, Lukasz Kaiser, and Illia Polosukhin. Attention Is All You Need. (Nips), 2017.

[32] Tomas Mikolov, Kai Chen, Greg Corrado, and Jeffrey Dean. Distributed Respresentation of Words and Pharses and their Compositionality. CrossRef Listing of Deleted DOIs, 2000.

[33] Jeffrey Pennington, Richard Socher, and Christopher Manning. Glove: Global Vectors for Word Representation. Proceedings of the 2014 Conference on Empirical Methods in Natural Language Processing (EMNLP), pages 1532–1543, 2014.

[34] Mark Stevenson and Yikun Guo. Disambiguation in the biomedical domain: The role of ambiguity type. Journal of Biomedical Informatics, 43(6):972–981, 2010.

[35] Matthew E. Peters, Mark Neumann, Mohit Iyyer, Matt Gardner, Christopher Clark, Kenton Lee, and Luke Zettlemoyer. Deep contextualized word representations. 2018.

[36] Jacob Devlin, Ming-Wei Chang, Kenton Lee, and Kristina Toutanova. BERT: Pre-training of Deep Bidirectional Transformers for Language Understanding. 2018.

[37] Chenguang Zhu, Michael Zeng, and Xuedong Huang. Sdnet: Contextualized attention-based deep network for conversational question answering. CoRR, abs/1812.03593, 2018.

[38] Alec Radford. Improving language understanding by generative pre-training. 2018.

[39] Shimi Salant and Jonathan Berant. Contextualized word representations for reading comprehension. In Proceedings of the 2018 Conference of the North American Chapter of the Association for Computational Linguistics: Human Language Technologies, Volume 2 (Short Papers), pages 554–559, New Orleans, Louisiana, 2018. Association for Computational Linguistics.

[40] Yanshan Wang, Sijia Liu, Naveed Afzal, Majid Rastegar-Mojarad, Liwei Wang, Feichen Shen, Paul Kingsbury, and Hongfang Liu. A comparison of word embeddings for the biomedical natural language processing. Journal of Biomedical Informatics, 87:12–20, 2018.

[41] Cédric Colas, Olivier Sigaud, and Pierre-Yves Oudeyer. How Many Random Seeds? Statistical Power Analysis in Deep Reinforcement Learning Experiments. pages 1–20, 2018.

[42] R. Kavuluru, A. Rios, and T. Tran. Extracting drug-drug interactions with word and character-level recurrent neural networks. In 2017 IEEE International Conference on Healthcare Informatics (ICHI), pages 5–12, Aug 2017.

[43] Chung Chi Huang and Zhiyong Lu. Community challenges in biomedical text mining over 10 years: Success, failure and the future. Briefings in Bioinformatics, 17(1):132–144, 2016.

[44] Eugene Charniak and Mark Johnson. Coarse-to-fine n-best parsing and maxent discriminative reranking. In Proceedings of the 43rd Annual Meeting on Association for Computational Linguistics, ACL’05, pages 173–180, Stroudsburg, PA, USA, 2005. Association for Computational Linguistics.

[45] Marie-Catherine De Marneffe, Bill MacCartney, and Christopher D. Manning. Generating typed dependency parses from phrase structure parses. Proceedings of the 5th International Conference on Language Resources and Evaluation (LREC 2006), pages 449–454, 2006.

[46] Sampo Pyysalo, Filip Ginter, Hans Moen, Tapio Salakoski, and Ananiadou Sophia. Distributional Semantics Resources for Biomedical Text Processing. Languages in Biology and Medicine, 2013.

[47] Shu Zhang, Dequan Zheng, Xinchen Hu, and Ming Yang. (2015) Bidirectional Long Short-Term Memory Networks for Relation Classification. pages 73–78, 2015.

[48] Mike Schuster and Kuldip K. Paliwal. Bidirectional recurrent neural networks. IEEE Trans. Signal Processing, 45:2673–2681, 1997.

[49] Zhiheng Huang, Wei Xu, and Kai Yu. Bidirectional LSTM-CRF Models for Sequence Tagging. 2015.

[50] Xuezhe Ma and Eduard Hovy. End-to-end Sequence Labeling via Bi-directional LSTM-CNNs-CRF. 2016.

[51] Yoon Kim. Convolutional neural networks for sentence classification. In Proceedings of the 2014 Conference on Empirical Methods in Natural Language Processing, EMNLP 2014, October 25-29, 2014, Doha, Qatar, A meeting of SIGDAT, a Special Interest Group of the ACL, pages 1746–1751, 2014.

[52] Wenpeng Yin, Katharina Kann, Mo Yu, and Hinrich Schütze. Comparative study of cnn and rnn for natural language processing. CoRR, abs/1702.01923, 2017.

[53] Zhenchao Jiang, Lishuang Li, Degen Huang, and Liuke Jin. Training word embeddings for deep learning in biomedical text mining tasks. In Proceedings - 2015 IEEE International Conference on Bioinformatics and Biomedicine, BIBM 2015, 2015.

[54] Hamed Zamani and W. Bruce Croft. Relevance-based Word Embedding. 2017.

[55] John S Bridle. Probabilistic Interpretation of Feedforward Classification Network Outputs, with Relationships to Statistical Pattern Recognition. In Françoise Fogelman Soulié and Jeanny Hérault, editors, Neurocomputing, pages 227–236, Berlin, Heidelberg, 1990. Springer Berlin Heidelberg.

[56] Adam Paszke, Sam Gross, Soumith Chintala, Gregory Chanan, Edward Yang, Zachary DeVito, Zeming Lin, Alban Desmaison, Luca Antiga, and Adam Lerer. Automatic differentiation in pytorch. In NIPS-W, 2017.

[57] Sebastian Ruder. Neural Transfer Learning for Natural Language Processing. PhD thesis, National University of Ireland, Galway, 2019.

[58] Matthew Peters, Mark Neumann, Luke Zettlemoyer, and Wen-tau Yih. Dissecting contextual word embeddings: Architecture and representation. In Proceedings of the 2018 Conference on Empirical Methods in Natural Language Processing, pages 1499–1509, Brussels, Belgium, October-November 2018. Association for Computational Linguistics.

[59] Kevin Clark, Minh-Thang Luong, Christopher D. Manning, and Quoc Le. Semi-supervised sequence modeling with cross-view training. In Proceedings of the 2018 Conference on Empirical Methods in Natural Language Processing, pages 1914–1925, Brussels, Belgium, October-November 2018. Association for Computational Linguistics.

